# Individual differences in the boldness of female zebrafish are associated with alterations in serotonin function

**DOI:** 10.1101/2024.02.13.580160

**Authors:** Fatemeh Beigloo, Cameron J. Davidson, Joseph Gjonaj, Shane A. Perrine, Justin W. Kenney

## Abstract

One of the most prevalent axes of behavioral variation in both humans and animals is risk taking, where individuals that are more willing to take risk are characterized as bold while those that are more reserved as shy. Brain monoamines (i.e., serotonin, dopamine, and norepinephrine) have been found to play a role in a variety of behaviors related to risk taking. Genetic variation related to monoamine function have also been linked to personality in both humans and animals. Using zebrafish, we investigated the relationship between monoamine function and boldness behavior during exploration of a novel tank. We found a sex-specific correlation between serotonin metabolism (5-HIAA:5-HT ratio) and boldness that was limited to female animals; there were no relationships between boldness and dopamine or norepinephrine. To probe differences in serotonergic function, we administered a serotonin reuptake inhibitor, escitalopram, to bold and shy fish, and assessed their exploratory behavior. We found that escitalopram had opposing effects on thigmotaxis in female animals with bold fish spending more time near the center of the tank and shy fish spent more time near the periphery. Taken together, our findings suggest that variation in serotonergic function makes sex-specific contributions to individual differences in risk taking behavior.

## Introduction

One of the most important and widely studied axes of behavioral variation is risk taking, where individuals that are more likely to take risks are characterized as bold, and those that are less likely as shy (Budaev and Brown, 2011; Réale et al., 2007; Wilson et al., 1994). Such individual differences in behavior are prevalent throughout the animal kingdom and are thought to contribute to evolutionary fitness (Dingemanse and Wolf, 2013; Réale et al., 2007). For example, individual differences in risk taking has been found to affect dispersal in great tits (Dingemanse et al., 2003) and migration in roach fish (Chapman et al., 2011). Furthermore, a recent meta-analysis found that higher risk taking in wild populations is associated with better survival (Moiron et al., 2020). In humans, increased risk taking in males is thought to contribute to their higher mortality compared to females (Kruger and Nesse, 2006; Wilson and Daly, 1985). However, despite our increased appreciation for the consequences of risk taking, we lack a detailed understanding of its biological basis.

One known contributor to behavioral variation is differences in the regulation of brain states by neuromodulators, like monoamines. Monoamines (i.e., dopamine, norepinephrine, and serotonin [5-hydroxytryptamine, 5-HT]), are a class of neuromodulators that are released from a small number of neuronal nuclei, but have extensive projections throughout the brain and regulate activity of many areas related to behavior (Jacobs and Azmitia, 1992). For example, serotonin has long been known to be involved behaviors related to boldness like aggression and risk assessment (Homberg and Lesch, 2011; Lucki, 1998). Similarly, dopamine has also been implicated in risk taking due, in part, to its role in modulating reward salience (Schultz, 2010). Accordingly, specific genetic alleles that contribute to serotonergic and dopaminergic function are associated with individual differences in risk taking in humans and animals, albeit with some inconsistency (Montag and Reuter, 2014).

Understanding the biological basis for individual differences benefits from the use of model organisms where we have greater control over conditions and access to a wide variety of molecular, pharmacological, and genetic tools. Zebrafish have recently gained popularity as a model organism for behavioral neurobiology due to their low cost, ease of genetic manipulation, extensive behavioral repertoire, and biological similarity to mammals (Gerlai, 2023; Kenney, 2020). Risk taking and boldness behavior in adult zebrafish has been characterized using a simple task called the novel tank test where animals explore a new environment. Risky behavior in this context consists of exploring more of the tank and spending time in particular parts of the tank, like the top, that would expose animals to greater risk of predation in the wild (Gerlai, 2020; Luca and Gerlai, 2012; Spence et al., 2006). Here, we tested the hypothesis that there is a relationship between boldness in a novel tank and monoamine function in the brain of zebrafish.

## Results

### Sex differences in behavior and brain chemistry

To assess the relationship between monoamine levels in the brain and exploratory behavior, we exposed fish to a novel tank on three consecutive days. Two days later (day 5), brains were dissected, and monoamine and metabolite levels were measured using HPLC (Figure 1A). Initially, we determined if there were sex differences in behavior and brain chemistry that would behoove us to analyze male and female fish separately (Figure 1B-D). Four exploratory behaviors were measured during exploration of the novel tank: two predator avoidance behaviors (bottom dwelling as measured by bottom distance and thigmotaxis as measured by center distance), activity levels (distance travelled) and how much of the tank fish explored (Figure 1B). Mixed two-way ANOVAs (sex × day) were used to assess significance. For bottom distance, there was a small effect of sex (*P* = 0.044, η^2^ = 0.029) where female fish spent more time near the top of the tank than males. There was no effect of day (*P* = 0.44) or an interaction between sex and day (*P* = 0.75). Center distance was not affected by sex (*P* = 0.25), but there was a medium sized effect of day (*P* < 0.0001, η^2^ = 0.11) and a small interaction (*P* = 0.034, η^2^ = 0.031). Both male and female fish increased their distance from the center over time, and on the first day female fish spent more time closer to the center than male fish. Distance travelled also changed across days and differed between the sexes: there was a large effect of sex (*P* < 0.001, η^2^ = 0.16), with male fish swimming further than female fish, and a small effect of day (*P* < 0.001, η^2^ = 0.032) such that activity decreased over time. There was no sex by day interaction (*P* = 0.51). Finally, there was a medium sized effect of sex on percent explored (*P* < 0.001, η^2^ = 0.082) where male fish explored more than female fish. There was neither an effect of day (*P* = 0.50) nor a sex by day interaction (*P* = 0.59) on percent explored.

**Figure 1.**
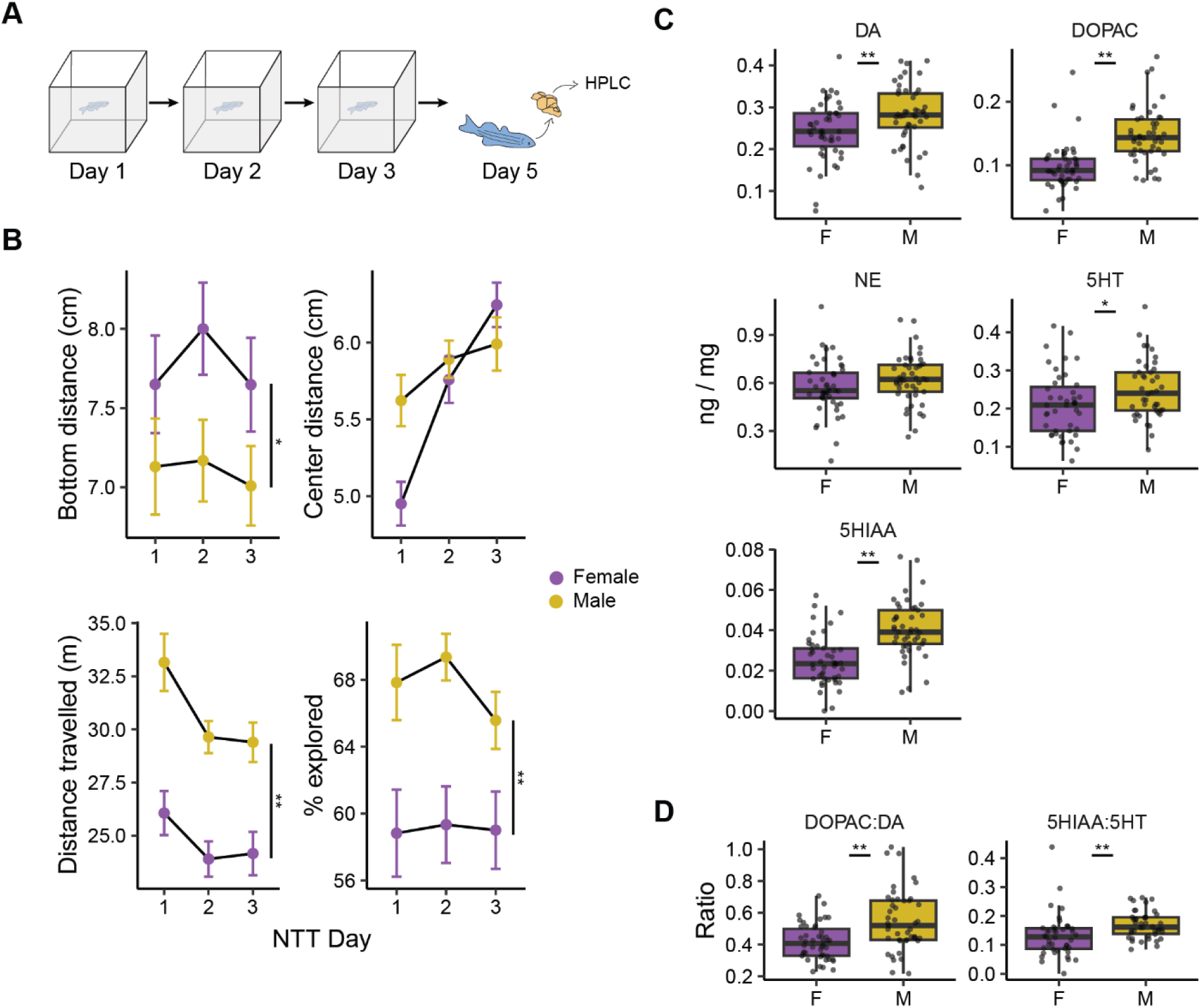
Sex differences in behavior and brain chemistry. A) Experimental design for assessing the relationship between exploration of a novel tank and brain chemistry. Image modified from Rajput et al (2022). B) The effect of sex and day on four exploratory behaviors in the novel tank. * - *P* < 0.05, ** - *P* < 0.001 indicate a main effect of sex from a sex × day ANOVA. Data are mean ± SEM. C) The effect of sex on monoamine and metabolite levels. D) The effect of sex on the ratio of monoamines to their metabolites. Boxplot center line is the median, hinges are interquartile ranges, and whiskers are the hinge ± 1.5 times the interquartile range. * - *P* < 0.05, ** - *P* < 0.01 from independent sample t-tests; n’s = 45-46.

We also determined if there were sex differences in the brain levels of monoamines or their metabolites, 5-HIAA (5-hydroxyindoleacetic acid) and DOPAC (3,4-dihydroxyphenylacetic acid) (Figure 1C). We found a medium sized effect of sex such that male fish had higher levels of dopamine (*P* = 0.0082, d = 0.57) and serotonin (*P* = 0.032, d = 0.47), and large effect on the metabolites DOPAC (*P* < 0.001, d = 1.23) and 5-HIAA (*P* < 0.001, d = 1.19). Levels of NE did not differ between the sexes (*P* = 0.12). We also assessed the turnover of dopamine and serotonin by calculating the DOPAC:DA and 5-HIAA:5-HT ratios (Figure 1D). We found that male fish had higher turnover than female fish for both neurotransmitters (DOPAC:DA: *P* = 0.0001, d = 0.87; 5-HIAA:5-HT: *P* = 0.005, d = 0.62). Taken together with the behavioral data, the sex differences we observed suggest that female and male fish should be analyzed separately.

### Correlations between boldness and brain chemistry

To assess the relationship between brain monoamine levels and boldness, we calculated a boldness index for each animal that combined the z-scores of bottom distance and percent explored. This is based on a previous work where we assessed behavior from over 400 animals and found that boldness in the novel tank test was comprised of a combination of willingness to explore and proximity to the top of the tank (Rajput et al., 2022). We then determined if monoamine and metabolite levels correlated with the boldness index on each day of testing (Figure 2A). Behavior on the first day of the novel tank was correlated with the 5-HIAA:5-HT ratio in female, but not male, zebrafish (Female: r = 0.37, *P* = 0.016; Male: r = -0.10, *P* = 0.53). In female zebrafish, two animals had high levels of 5-HIAA:5-HT that might unduly influence the correlation (Figure 2B), so we also calculated the non-parametric Spearman’s ρ that is less susceptible to outliers, with essentially the same result (ρ = 0.36, *P* = 0.02). There were also three significant correlations between boldness and brain chemistry in male fish, but none on the first day (Figure 2A): on the second and third days higher 5-HIAA levels were associated with less boldness (Day 2: r = -0.29, *P* = 0.050; Day 3: r = -0.31, *P* = 0.040). On the third day, the DOPAC:DA ratio correlated with boldness in male fish (r = 0.30, *P* = 0.043).

**Figure 2.**
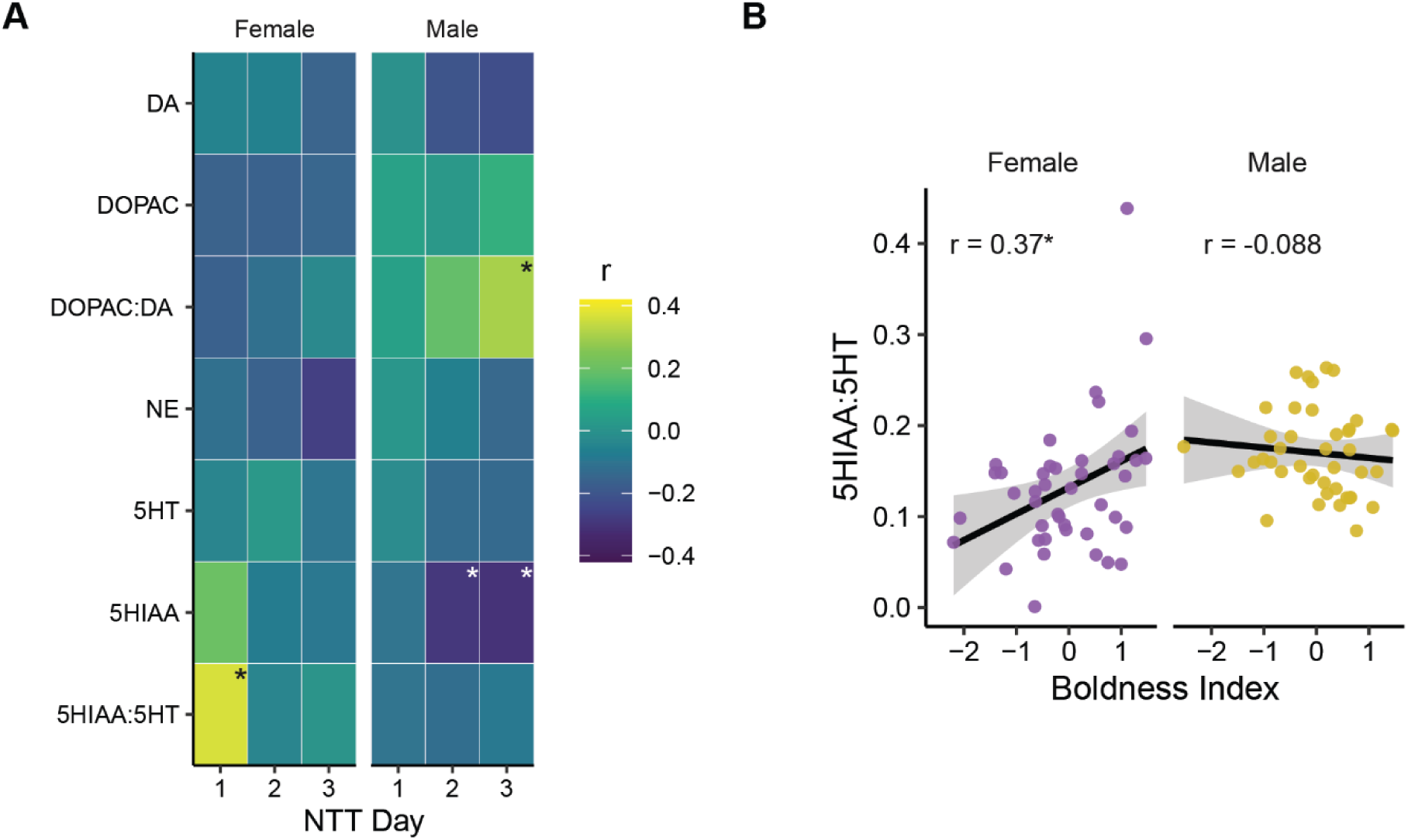
Correlations between boldness index and brain chemistry. A) Pearson’s correlations between the boldness index on each day of testing in the novel tank and monoamines, their metabolites, or the ratio of metabolites to monoamines. B) Correlation between the 5-HIAA:5-HT ratio and boldness index on day one of testing in the novel tank. * - *P* < 0.05.

We also calculated correlations between individual behaviors and brain chemistry (Figure S1). There were two other significant correlations in females, both for distance travelled (Figure S1C): on day 1 we found a positive relationship with the 5-HIAA:5-HT ratio (r = 0.32, *P* = 0.036; Figure S1C) and on day 2 a negative relationship with norepinephrine (r = -0.31, *P* = 0.035). For male fish, there were no significant correlations on day 1 (Figure S1). On days two and three there were several negative relationships with bottom distance (Figure S1A): On day 2, with the 5-HIAA:5-HT ratio (r = -0.31, *P* = 0.048) and with 5-HIAA on both days 2 (r = -0.45, *P* = 0.0021) and 3 (r = -0.40, *P* = 0.0068). The only other significant correlations in males were with distance travelled on day 2 (Figure S1C) where there was a positive relationship with the 5- HIAA:5-HT ratio (r = 0.47, *P* = 0.0015) and a negative relationship with 5-HT (r = -0.32, *P* = 0.042).

### Blocking serotonin reuptake has different effects in bold and shy female zebrafish

The correlation between boldness and the 5-HIAA:5-HT ratio in female fish on day 1 suggests a role for the serotonergic system in this behavior. Based on this, we hypothesized that bold and shy female fish would respond differently to a drug that targets the serotonergic system. To test this hypothesis, we administered escitalopram, a serotonin reuptake inhibitor (Sánchez et al., 2004), to bold and shy fish prior to exploration of a tank (Figure 3A). To identify bold and shy fish, we exposed animals to the novel tank and did a median split on the boldness index within sex (Figure 3B). A day later, fish were administered either vehicle or 1 mg/kg of escitalopram 30 minutes before being placed back into the tank. We assessed four behaviors on the second day (bottom distance, center distance, distance travelled, and percent tank explored) using a 2 × 2 (drug × boldness) ANOVA design.

**Figure 3.**
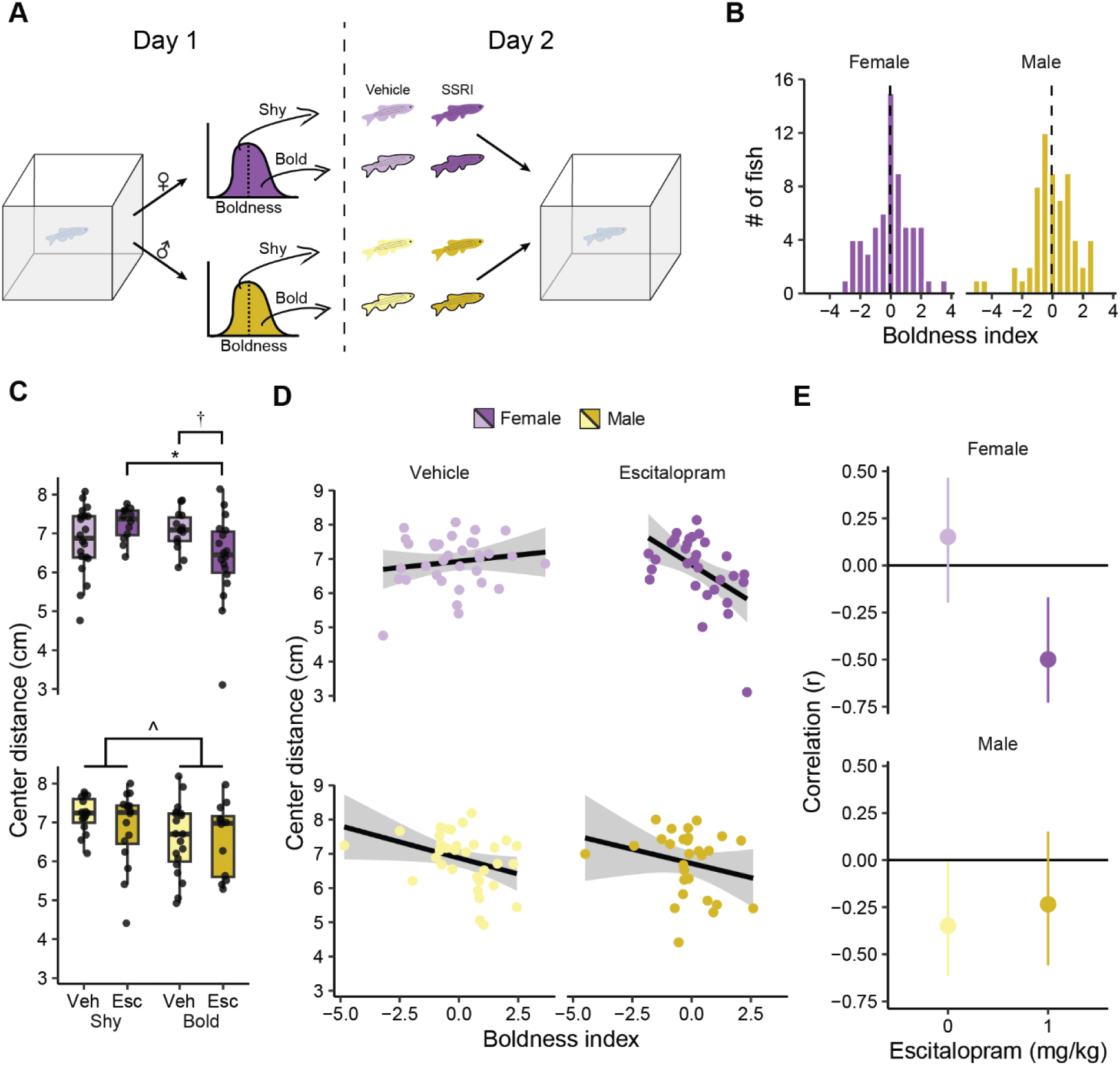
The effect of escitalopram on the exploratory behavior of bold and shy fish. A) Fish were separated into bold and shy animals based on a median split of their boldness index during an initial exposure to the tank (day 1). On day 2, fish were given either vehicle or 1 mg/kg of escitalopram prior to being placed back into the novel tank. B) Histogram of boldness index on day 1 of exposure to the novel tank. The dashed line is the median. C) Center distance on day 2 in female (top) and male (bottom) animals. Boxplot center is the median, hinges are interquartile ranges, and whiskers are the hinge ± 1.5 times the interquartile range. D) Scatterplots and regression lines for boldness index on day 1 and center distance on day 2. E) Pearson’s correlations and 95% confidence intervals (error bars) that correspond to scatterplots in part D. † - *P* < 0.10, * - *P* < 0.05 from FDR corrected post-hoc t-tests. ^ - *P* < 0.05 from two-way ANOVA; shy female vehicle, n = 20; shy female escitalopram, n = 12; bold female vehicle, n = 14; bold female escitalopram, n = 18; shy male vehicle, n = 15; shy male escitalopram, n = 16; bold male vehicle, n = 19; bold male escitalopram, n = 12.

The effect of escitalopram on center distance in female fish depended on whether animals were categorized as bold or shy (Figure 3C, top) as evidenced by a medium sized interaction between boldness and drug (*P* = 0.0093, η^2^ = 0.10): Shy animals given the drug increased their center distance whereas bold fish decreased their center distance. Follow-up FDR corrected pair-wise t-tests found a difference between bold and shy animals given escitalopram (*P* = 0.038) and a trend towards a difference between bold fish given vehicle and drug (*P* = 0.062). There were no main effects of boldness (*P* = 0.16) or drug (*P* = 0.48). In a separate analysis, male fish demonstrated no effect of the drug (*P* = 0.45) and a medium sized effect of boldness (*P* = 0.047, η^2^ = 0.071) where animals that were bold on day one spent more time closer to the center of the tank on day 2. Different from the females, no such interaction between drug and boldness was seen (*P* = 0.43; Figure 3C, bottom).

We also determined if boldness on day 1 correlated with center distance on day 2 and whether this was affected by escitalopram treatment (Figures 3D & E). In vehicle treated female fish, there was no correlation between boldness on day 1 and center distance on day 2 (r = 0.15, *P* = 0.39) whereas there was a moderate negative correlation in those given escitalopram (r = -0.50, *P* = 0.005). The ninety-five percent confidence intervals for vehicle or escitalopram treated female fish slightly overlapped (Figure 3E, top). In male fish, there was a weak negative correlation between boldness index on day 1 and center distance on day 2 in vehicle treated animals (r = -0.35, *P* = 0.044), and no correlation in those given escitalopram (r = -0.23, *P* = 0.23). The ninety-five percent confidence intervals for vehicle and escitalopram treated males largely overlapped (Figure 3E, bottom).

There were no interactions between boldness and drug for other behavioral parameters (Figure S2). For bottom distance (Figure S2A), we found a trend towards a small effect of boldness in female fish (*P* = 0.052, η^2^ = 0.058) where bolder fish spend more time near the top of the tank. There was no effect of drug treatment (*P* = 0.99) or interaction with boldness (*P* = 0.27). In male fish, bottom distance was not affected by day 1 boldness (*P* = 0.12) or drug (*P* = 0.25) and there was no interaction (*P* = 0.45). Locomotor activity (Figure S2B) was not affected by escitalopram in female fish (*P* = 0.18). In male fish, there was a trend towards a small effect of drug (*P* = 0.092, η^2^ = 0.048) where fish given escitalopram swam less. There were no effects of boldness or an interaction in either female or male fish on locomotor activity (Female: boldness: *P* = 0.81, interaction, *P* = 0.31. Male: boldness: *P* = 0.88, interaction: *P* = 0.74). The percent of the tank explored on day 2 (Figure S2C) was not affected by boldness on day 1 in female fish (*P* = 0.45), but there was a trend towards a small effect of escitalopram (*P* = 0.072, η^2^ = 0.052) where fish given the drug explored less than those given vehicle. There was no interaction between boldness and drug treatment (*P* = 0.34). In male fish, there was a medium sized effect of boldness on percent explored (*P* = 0.022, η^2^ = 0.73) such that bold fish explored more than shy fish (Figure 3F, bottom). There was no effect of drug (*P* = 0.30) or an interaction between boldness and drug (*P* = 0.23).

We also determined whether escitalopram affected correlations between bottom distance, distance travelled, and percent explored on day 2 and boldness index on day 1 (Figure S3). In vehicle treated female fish there was a moderate positive correlation between day 1 boldness index and day 2 bottom distance (r = 0.58, *P* < 0.001; Figure S3A, top) and weak correlation between boldness index on day 1 and percent of tank explored on day 2 (r = 0.34, *P* = 0.046; Figure S3C, top). For bottom distance, the 95-percent confidence intervals between vehicle and escitalopram treated female fish overlapped slightly (Figure S3A, top). There was no correlation between boldness index on day 1 and distance travelled in female fish (Figure S3B, top). In male fish, there were no significant correlations between boldness index on day 1 and behavior on day 2, and all 95-percent confidence intervals in vehicle and drug treated fish had large overlaps (Figure S3, bottom).

## Discussion

Our findings identify the serotonergic system as playing a sex-specific role in individual differences in the boldness behavior of zebrafish. This conclusion is based on several findings from the present study: (1) we found a correlation between boldness during an initial exposure to a novel tank and serotonin turnover in the brain that was limited to female fish (Figure 2). (2) Elevating serotonin at the synapse of female fish using a serotonin reuptake inhibitor, escitalopram, resulted in decreased thigmotaxis in bold animals and increased thigmotaxis in shy animals (Figure 3C). (3) Escitalopram treatment affected correlations between boldness during an initial exposure to a novel tank and behavior on a second exposure in female, but not male, zebrafish (Figures 3D-E; Figure S3). Taken together, our data suggests an important role for serotonin in the emergence of individual differences in risk taking during exploration of a novel environment, particularly in female animals.

The serotonergic system of zebrafish is characterized by several populations of neurons, some of which resemble those seen in mammals, like the raphe nuclei, and others that appear to be unique to fish, like clusters in the hypothalamus and pretectum (reviewed in Lillesaar, 2011). Despite these differences in neuroanatomy, the behavioral functions of the serotonergic system are remarkably conserved, implicated in aggression, fear, and anxiety in both zebrafish and mammals (Herculano and Maximino, 2014; Lillesaar, 2011; Lucki, 1998; Winberg and Thörnqvist, 2016). This may be due to the fact that, like in mammals, many long range serotonergic projections in zebrafish originate from the raphe nuclei (Lillesaar et al., 2009) and those projections missing in fish often contain local populations of serotonergic neurons that may subserve a similar function (Lillesaar, 2011). Given this high level of conservation, findings from the present study are likely to provide insight into serotonergic function across species.

The most striking finding from this study is that bold and shy female zebrafish respond in opposite ways to treatment with escitalopram: drug treated bold fish spent more time in the center of the tank whereas shy fish spent more time in the periphery (Figure 3). This echoes work in mammals where individual differences in the open field activity of rats are associated with opposing effects of different serotonergic drugs on time to enter the open arms of an elevated T-maze (Verheij et al., 2009), and in humans, where the acute effects of modifying serotonergic function on moral judgment and social emotions depends on personality traits (Crockett et al., 2010; Kanen et al., 2021). However, the question remains: how does elevating serotonin at the synapse result in differential behavioral responses in bold and shy animals? One likely culprit is differences in serotonergic receptor expression and/or localization throughout the brain. Serotonin receptors are highly conserved and are divided into four classes based on their signaling mechanisms (Nichols and Nichols, 2008). Most receptors are G-protein coupled, except for 5-HT_3_, which is a cation permeable ligand gated ion channel. Further complicating the picture in zebrafish is that, due to the fish specific genome duplication event that occurred ∼350 million years ago (Meyer and Van de Peer, 2005), many serotonin receptors have multiple isoforms unique to fish (Maximino et al., 2016; Norton et al., 2008). Nonetheless, it is known that modulation of different classes of serotonergic receptors have opposite influences on the exploratory behavior of zebrafish: 5-HT_1A_ and 5-HT_1B/D_ antagonists increase bottom distance whereas a 5-HT_2_ receptor antagonist decrease this behavior (Maximino et al., 2013; Nowicki et al., 2014). Thus, it is likely that variation in the expression of 5-HT receptors throughout the brain underlie individual differences in boldness.

We also found that the ratio of 5-HIAA:5-HT correlated with boldness in female zebrafish such that animals with a higher ratio were bolder than those with a lower ratio. This higher turnover could reflect differences in the synthesis or degradation of serotonin. Degradation of 5- HT to 5-HIAA is catalyzed by monoamine oxidase, for which zebrafish have only one gene instead of two as in mammals (Setini et al., 2005). In contrast, zebrafish have four genes that encode the rate-limiting enzyme for serotonin synthesis, tryptophan hydroxylase (*tph1a*, *tph1b*, *tph2*, and *tph3*), compared to two in mammals (Ren et al., 2013; Walther and Bader, 2003). It is likely that these three isoforms have different kinetics, however, only the kinetics of *tph3* (formerly known as *th2*) are currently known (Ren et al., 2013). The expression of these isoforms varies throughout the brain (Bellipanni et al., 2002; Lillesaar et al., 2007; Ren et al., 2013; Teraoka et al., 2004), and thus, it may be the case that the differences in the turnover of 5-HT in bold and shy fish may reflect distinct distributions of tyrosine hydroxylase isoforms.

Two prior studies have examined links between behavioral variation and monoamine levels in the brains of zebrafish. Tran and colleagues (2016) found that zebrafish with high locomotor activity had lower levels of brain serotonin, with no difference in dopamine. We also observed a negative correlation between distance travelled and serotonin, albeit only in male fish and only on the second day of exposure to the tank (Figure S1C). Another study found a correlation between serotonin and time spent in the top of a novel tank (Maximino et al., 2013), an effect we did not observe (Figure S1A). Several methodological differences may explain the discrepancies between the present study and prior work. In the present study, brain dissections occurred two days after the last behavioral manipulation so as to capture baseline levels of chemicals in the brain. In contrast, both prior studies collected samples immediately after exposure to a novel tank. Given that exposure to the novel tank test itself is known to elicit region specific elevations in serotonin (Maximino et al., 2013), a direct comparison of our findings with these studies is difficult. It may be the case that serotonin levels increased from baseline in some animals more than others in response to the behavioral manipulation. In addition, we found several sex differences in behavior and monoamine levels, with, for example, the relationship between locomotor activity and serotonin to be restricted to male fish. Prior studies did not indicate sex ratios or stratify results by sex (Maximino et al., 2013; Tran et al., 2016). Nonetheless, as a whole, these studies support the conclusion that serotonin, but not dopamine, contributes to individual differences in the exploratory behavior of zebrafish.

The link between serotonin levels in the brain and variation in boldness-related behaviors is not limited to zebrafish, but has also been found in other fish species, rodents, and invertebrates like crayfish. For example, similar to our findings, bold European sea bass and hatchery reared Atlantic cod have higher 5-HIAA:5-HT ratios than their shy counterparts (Alfonso et al., 2019; Meager et al., 2012). In male rats, animals that spend less time in the open arms of an elevated plus maze, and thus are considered more anxious or shy, have higher levels of serotonin in the amygdala, but not other brain regions like the hippocampus or striatum (Näslund et al., 2015). This brain region specificity in rats may be why we do not observe a similar relationship in male fish in our study since we used whole-brain homogenates. Similarly, in crayfish, serotonin, but not dopamine, levels were higher in animals that spend more time in darker parts of a maze than less anxiety-provoking light parts (Fossat et al., 2015). These findings across phyla speak to the high evolutionary conservation of the contribution of serotonin to individual behavioral variation despite wide variations in neuroanatomy (Backström and Winberg, 2017; Herculano and Maximino, 2014).

Prior work using escitalopram, or another serotonin reuptake inhibitor, fluoxetine, found that acute administration alters behavior in the novel tank test, increasing time spent near the top of the tank (Fontana et al., 2022; Lima-Maximino et al., 2020; Maximino et al., 2013; Sackerman et al., 2010). Here, however, we did not find that escitalopram increased bottom distance (Figure S2A). This is likely due to the fact that escitalopram was given prior to a second exposure to the tank, when the tank was no longer novel (Figure 3A). This suggests that novelty may be an important element that engages the serotonergic system. A link between novelty and serotonin is further supported by our finding that the correlation between boldness and serotonin turnover is only present during the initial exposure to the novel tank, but not subsequent days (Figure 2A). Interestingly, polymorphisms in serotonin transporter genes in humans are associated with novelty seeking, often in a sex-specific manner (Kazantseva et al., 2008; Vormfelde et al., 2006). Thus, it may be that novelty is important for engaging the serotonergic system in the context of risk taking and it may differ between sexes.

We found several differences between male and female fish: the link between serotonin and boldness was limited to female zebrafish, and females had lower levels of most neurotransmitters and metabolites (except for norepinephrine), and lower DOPAC:DA and 5-HIAA:5-HT ratios than males. Similarly, Dahlbom and colleagues (2012) found that females had lower 5-HIAA:5-HT and DOPAC:DA ratios in the forebrain, and lower 5-HT in the hindbrain, but elevated dopamine in the forebrain. The difference in dopamine levels between prior work and the present study may be due to our use of whole-brain homogenates compared to dividing the brain into forebrain and hindbrain. Indeed, regional heterogeneity in sex differences in monoamine levels has been reported in rodents (Bowman et al., 2009; De la Fuente et al., 2003; Liiver et al., 2023), suggesting that it is a widespread phenomenon. These sex differences may reflect differences in tryptophan and/or serotonin metabolism due to the regulation of related genes by sex hormones (Barth et al., 2015). How these sex differences in neurochemistry give rise to sexual dimorphism in behavior will likely require a closer examination of regional differences.

One of the limitations of the present study is the use of whole-brain homogenates for HPLC to identify monoamines and their metabolites. This was necessitated, in part, by the relatively small size of the zebrafish brain to ensure we obtained enough material for detection. Future work should explore potential regional differences for the contribution of the serotonergic system to zebrafish behavior in more detail by taking advantage of recent advances in tools for whole-brain mapping in adult zebrafish (Kenney et al., 2021). Another limitation to the present work was the use of escitalopram, a serotonin reuptake inhibitor, which is expected to increase serotonin at the synapse throughout the brain. We used such a broad approach expecting that it was likely to reveal differences that may be attributable to multiple aspects of serotonergic function. A fruitful avenue for future work might be to use receptor specific drugs in bold and shy animals or to knock out receptor genes using recent advances in CRISPR/Cas9 (Kroll et al., 2021).

The suggested involvement of serotonin in personality and individual behavioral differences goes back nearly half a century. However, a mechanistic understanding of how serotonin contributes to behavioral variation has remained elusive. By uncovering a role for serotonin in the individual differences of boldness in zebrafish, the present study lays the foundation for using the rapidly growing toolbox available to zebrafish researchers to shed light on how variation in serotonergic function gives rise to individual differences in behavior.

## Methods

### Subjects

Subjects were female and male zebrafish of the WIK strain within two generations of animals originally from the Zebrafish International Resource Center (ZIRC, catalog ID: ZL84) at the University of Oregon. Fish were 16-35 weeks post fertilization and raised at Wayne State University. Housing was in high density racks under standard conditions (water temperature: 27.5 ± 0.5 ⁰C, salinity: 500 ± 10 µS, and pH = 7.4 ± 0.2) with a 14:10 light:dark cycle (lights on at 8:00AM). Fish were fed twice daily, in the morning with a dry feed (Gemma 300, Skretting, Westbrook, ME, USA) and in the afternoon with brine shrimp (*Artemia salina*, Brine Shrimp Direct, Ogden, UT, USA).

The sex of fish was determined using three secondary sex characteristics: shape, color, and presence of pectoral fin tubercles (McMillan et al., 2015). Sex was confirmed following euthanasia by the presence or absence of eggs. All procedures were approved by the Wayne State University Institutional Animal Care and Use Committee.

### Novel tank test

One week prior to behavioral manipulations, animals were placed into 2 L tanks with two female/male pairs per tank. Tanks were divided in half with a transparent divider with one pair per side. This arrangement allowed us to keep track of fish identity over days without social isolation or tagging. One hour before exposure to the novel tank, animals were removed from their housing racks and brought to the behavioral room to acclimate. Following testing, fish remained in the behavioral room for at least 30 minutes before being returned to housing racks. Fish were individually placed into experimental tanks for 6 minutes while videos were recorded. Water was replaced between animals to prevent the buildup of chemical cues that may be released by fish during the test.

Novel tanks were five-sided (15 ˣ 15 ˣ 15 cm) and made from frosted acrylic (TAP Plastics, Stockton, CA, USA). Tanks were filled to a height of 12 cm with 2.5 L of fish facility water and placed in a white plasticore enclosure to prevent disturbance by external stimuli and to diffuse light. The tanks were open on the top, above which D435 Intel RealSense^TM^ cameras (Intel, Santa Clar, CA, USA) were placed for recording. Cameras were mounted 20 cm above the tanks and connected to Linux workstations using high-speed USB cables (NTC distributing, Santa Clara, CA, USA) for recording.

### Animal tracking

D435 cameras have color and depth video streams that create three-dimensional videos (Kuroda, 2018; Rajput et al., 2022). The color video was extracted from each file and five points along the fish were tracked using DeepLabCut (Mathis et al., 2018). Three-dimensional swim traces were created by overlaying the tracking with the depth video using custom written Python code. We extracted four behavioral parameters as previously described (Rajput et al., 2022): (1) distance from bottom where a plane was fit to the bottom of the tank and the distance from the fish to the plane was calculated, (2) center distance where the distance of the fish to a line down the center of the tank was calculated, (3) distance travelled, and (4) percent of tank explored where the tank was divided into one thousand evenly spaced voxels and the proportion of voxels visited was calculated. Prior to calculations, swim traces were smoothed using a Savitzky-Golay filter with a length of seven frames and an order of three (Press and Teukolsky, 1990).

### HPLC

HPLC analysis was conducted as previously described (Davidson et al., 2022; Gheidi et al., 2023). Following euthanasia in ice cold water, fish brains were rapidly dissected, flash frozen on dry ice and stored at -80 ⁰C. Whole brains were then thawed, weighed, and homogenized by sonication (XL-2000, Misonix, Durham, NC, USA) for 3-5 seconds in 50 µL of 0.2 M perchloric acid. Homogenate was visually inspected for uniform tissue suspension and then centrifuged at 10,000 g for 10 min at 4 ⁰C, and supernatant carefully retained. A 20 µL aliquot of each sample was placed in a tube and fed into an autosampler of a Dionex Ultimate 3000 HPLC system at 5 ⁰C (Thermo Fisher Scientific, Waltham, MA, USA). External standards for DA, 5-HT, NE, DOPAC, and 5-HIAA were prepared via serial dilution (0.1 to 10 ng) including a blank of perchloric acid. Standards were processed in ascending concentrations and run in parallel and duplicate. Dopamine metabolites 3-methoxytyramine and homovanillic acid were not assessed because they could not be reliably detected.

Standards and samples were injected (10 µL) by an autoinjector using an acetonitrile based MD-TM mobile phase (10% acetonitrile: Thermo Fisher Scientific). HPLC flow rate was 0.6 mL / min. Samples passed through a reverse-phase column (Hypersil^TM^ BDS C18 column, Thermo Fisher Scientifc) at 25 ⁰C, a guard cell (ESA guard cell, model 2050) set to 300 mV, and a column guard (2.1/3.0 mm ID, Thermo Fisher Scientific). For electrochemical detection, we used an ultra-analytical dual-electrode cell (ESA, model 5011A) with a reference electrode set to -175 mV and a working electrode at 350 mV (gain of 100 µA for both). Detector values were captured using Chromileon 7 software (Dionex, Thermo Fisher Scientific) to quantify peak height, representing monoamine and metabolite levels. A detection threshold of three times the perchloric acid background was set. Samples that did not reach threshold were excluded from analysis. Standard curves for each chemical were generated to calculate levels and verify instrument stability (R^2^ ≥ 0.97). Levels of monoamines are expressed as ng per mg of wet brain weight.

### Drug treatment

Escitalopram oxalate (Sigma-Aldrich, St. Louis, MO, USA) was administered using a gelatin-based feed at a dose of 1 mg of drug per kg of body weight as previously described (Ochocki and Kenney, 2023). The gelatin feed consisted of 12 % w/v gelatin (Sigma-Aldrich), 4 % w/v spirulina (Argent Aquaculture, Redmon, WA, USA) and brine shrimp extract. The extract was made by suspending 250 mg / mL of mikro fine brine shrimp (Brine Shrimp Direct) in water followed by 1 hour of stirring. The suspension was then centrifuged twice at 12,500 g, keeping the supernatant each time, and then diluted in two volumes of water before addition to gelatin feed mixture. Escitalopram stock (10 mg / mL) or vehicle (water) was added prior to warming the solution to 45 ⁰C. After warming, drug or vehicle containing solution was pipetted into individually sized morsels for feeding at 1% body weight. Gelatin was allowed to set at -20 ⁰C for at least 20 minutes prior to feeding.

Fish were given two days to acclimate to the gelatin feed prior to drug administration. On each day of the experiment, animals were removed from the housing racks to the behavioral room for one hour before being given a non-dosed gelatin feed in lieu of their morning feed. Fish were briefly isolated by placement of transparent barriers in the tanks for 2-5 minutes. Once feed was given, we recorded if fish ate the feed within five minutes. On day one of the novel tank test, all animals were given non-dosed feed 30 minutes prior to being placed in the novel tank. On the second day of testing, fish were given either vehicle or escitalopram containing feed 30 minutes prior to being placed in the novel tank.

### Statistical analysis

Data analysis was performed using version 4.3.0 of R (R Core Team, 2016). Graphs were made using ggplot2 (Wickham, 2015). Statistical analysis was done using either a 2 ˣ 3 (sex ˣ day) mixed ANOVA, 2 ˣ 2 (drug ˣ boldness) ANOVA, independent samples t-tests or Pearson’s/Spearman’s correlations as indicated. Interactions of interest from ANOVAs were followed up with pair-wise t-tests and corrected for multiple comparisons using the false discover rate (FDR) (Benjamini and Hochberg, 1995). Effect sizes for ANOVAs are reported as η^2^ and t-tests as Cohen’s d. The interpretation of effect sizes as small (0.01 < η^2^ < 0.06; 0.2 < d < 0.5), medium (0.06 ≤ η^2^ < 0.14; 0.5 ≤ d < 0.8), or large (η^2^ ≥ 0.14; d ≥ 0.8) are based on Cohen (1988).

## Acknowledgements

This work was funded by the National Institutes of Health (R35GM142566) to J.W.K. and to S.A.P. (R01DA042057 and R21DA052657).

## Notes

### Competing Interest Statement

The authors have declared no competing interest.

## References

Alfonso S, Sadoul B, Gesto M, Joassard L, Chatain B, Geffroy B, Bégout M-L. 2019. Coping styles in European sea bass: The link between boldness, stress response and neurogenesis. Physiology & Behavior 207:76–85. doi:10.1016/j.physbeh.2019.04.020

Backström T, Winberg S. 2017. Serotonin Coordinates Responses to Social Stress—What We Can Learn from Fish. Frontiers in Neuroscience 11.

Barth C, Villringer A, Sacher J. 2015. Sex hormones affect neurotransmitters and shape the adult female brain during hormonal transition periods. Frontiers in Neuroscience 9.

Bellipanni G, Rink E, Bally-Cuif L. 2002. Cloning of two tryptophan hydroxylase genes expressed in the diencephalon of the developing zebrafish brain. Mechanisms of Development 119:S215–S220. doi:10.1016/S0925-4773(03)00119-9

Benjamini Y, Hochberg Y. 1995. Controlling the False Discovery Rate: A Practical and Powerful Approach to Multiple Testing. Journal of the Royal Statistical Society: Series B (Methodological) 57:289–300. doi:10.1111/j.2517-6161.1995.tb02031.x

Bowman RE, Micik R, Gautreaux C, Fernandez L, Luine VN. 2009. Sex-dependent changes in anxiety, memory, and monoamines following one week of stress. Physiology & Behavior 97:21–29. doi:10.1016/j.physbeh.2009.01.012

Budaev S, Brown C. 2011. Personality Traits and BehaviourFish Cognition and Behavior. John Wiley & Sons, Ltd. pp. 135–165. doi:10.1002/9781444342536.ch7

Chapman BB, Hulthén K, Blomqvist DR, Hansson L-A, Nilsson J-Å, Brodersen J, Anders Nilsson P, Skov C, Brönmark C. 2011. To boldly go: individual differences in boldness influence migratory tendency. Ecology Letters 14:871–876. doi:10.1111/j.1461-0248.2011.01648.x

Cohen J. 1988. Statistical Power Analysis for the Behavioral Sciences, 2nd ed. New York: Routledge. doi:10.4324/9780203771587

Crockett MJ, Clark L, Hauser MD, Robbins TW. 2010. Serotonin selectively influences moral judgment and behavior through effects on harm aversion. Proceedings of the National Academy of Sciences 107:17433–17438. doi:10.1073/pnas.1009396107

Dahlbom SJ, Backström T, Lundstedt-Enkel K, Winberg S. 2012. Aggression and monoamines: Effects of sex and social rank in zebrafish (Danio rerio). Behavioural Brain Research 228:333–338. doi:10.1016/j.bbr.2011.12.011

Davidson CJ, Svenson DW, Hannigan JH, Perrine SA, Bowen SE. 2022. A novel preclinical model of environment-like combined benzene, toluene, ethylbenzene, and xylenes (BTEX) exposure: Behavioral and neurochemical findings. Neurotoxicology and Teratology 91:107076. doi:10.1016/j.nt.2022.107076

De la Fuente M, Hernanz A, Medina S, Guayerbas N, Fernández B, Viveros MP. 2003. Characterization of monoaminergic systems in brain regions of prematurely ageing mice. Neurochemistry International 43:165–172. doi:10.1016/S0197-0186(02)00212-7

Dingemanse NJ, Both C, van Noordwijk AJ, Rutten AL, Drent PJ. 2003. Natal dispersal and personalities in great tits (Parus major). Proceedings of the Royal Society of London Series B: Biological Sciences 270:741–747. doi:10.1098/rspb.2002.2300

Dingemanse NJ, Wolf M. 2013. Between-individual differences in behavioural plasticity within populations: causes and consequences. Animal Behaviour, Special Issue: Behavioural Plasticity and Evolution 85:1031–1039. doi:10.1016/j.anbehav.2012.12.032

Fontana BD, Alnassar N, Parker MO. 2022. The zebrafish (Danio rerio) anxiety test battery: comparison of behavioral responses in the novel tank diving and light–dark tasks following exposure to anxiogenic and anxiolytic compounds. Psychopharmacology 239:287–296. doi:10.1007/s00213-021-05990-w

Fossat P, Bacqué-Cazenave J, De Deurwaerdère P, Cattaert D, Delbecque J-P. 2015. Serotonin, but not dopamine, controls the stress response and anxiety-like behavior in the crayfish Procambarus clarkii. Journal of Experimental Biology 218:2745–2752. doi:10.1242/jeb.120550

Gerlai R. 2023. Zebrafish (Danio rerio): A newcomer with great promise in behavioral neuroscience. Neuroscience & Biobehavioral Reviews 144:104978. doi:10.1016/j.neubiorev.2022.104978

Gerlai RT. 2020. Chapter 10 - Fear responses and antipredatory behavior of zebrafish: a translational perspective In: Gerlai RT, editor. Behavioral and Neural Genetics of Zebrafish. Academic Press. pp. 155–171. doi:10.1016/B978-0-12-817528-6.00010-3

Gheidi A, Davidson CJ, Simpson SC, Yahya MA, Sadik N, Mascarin AT, Perrine SA. 2023. Norepinephrine depletion in the brain sex-dependently modulates aspects of spatial learning and memory in female and male rats. Psychopharmacology 240:2585–2595. doi:10.1007/s00213-023-06453-0

Herculano AM, Maximino C. 2014. Serotonergic modulation of zebrafish behavior: Towards a paradox. Progress in Neuro-Psychopharmacology and Biological Psychiatry, Special Issue: Zebrafish models of brain disorders 55:50–66. doi:10.1016/j.pnpbp.2014.03.008

Homberg JR, Lesch K-P. 2011. Looking on the Bright Side of Serotonin Transporter Gene Variation. Biological Psychiatry, Genes and Anxiety 69:513–519. doi:10.1016/j.biopsych.2010.09.024

Jacobs BL, Azmitia EC. 1992. Structure and function of the brain serotonin system. Physiological Reviews 72:165–229. doi:10.1152/physrev.1992.72.1.165

Kanen JW, Arntz FE, Yellowlees R, Cardinal RN, Price A, Christmas DM, Apergis-Schoute AM, Sahakian BJ, Robbins TW. 2021. Serotonin depletion amplifies distinct human social emotions as a function of individual differences in personality. Transl Psychiatry 11:1–12. doi:10.1038/s41398-020-00880-9

Kazantseva AV, Gaysina DA, Faskhutdinova GG, Noskova T, Malykh SB, Khusnutdinova EK. 2008. Polymorphisms of the serotonin transporter gene (5-HTTLPR, A/G SNP in 5-HTTLPR, and STin2 VNTR) and their relation to personality traits in healthy individuals from Russia. Psychiatric Genetics 18:167. doi:10.1097/YPG.0b013e328304deb8

Kenney JW. 2020. Associative and nonassociative learning in adult zebrafishBehavioral and Neural Genetics of Zebrafish. Elsevier. pp. 187–204. doi:10.1016/b978-0-12-817528-6.00012-7

Kenney JW, Steadman PE, Young O, Ting Shi M, Polanco M, Dubaishi S, Mueller T, Frankland PW. 2021. AZBA: A 3D Adult Zebrafish Brain Atlas for the Digital Age. bioRxiv 2021.05.04.442625. doi:10.1101/2021.05.04.442625

Kroll F, Powell GT, Ghosh M, Gestri G, Antinucci P, Hearn TJ, Tunbak H, Lim S, Dennis HW, Fernandez JM, Whitmore D, Dreosti E, Wilson SW, Hoffman EJ, Rihel J. 2021. A simple and effective F0 knockout method for rapid screening of behaviour and other complex phenotypes. eLife 10:e59683. doi:10.7554/eLife.59683

Kruger DJ, Nesse RM. 2006. An evolutionary life-history framework for understanding sex differences in human mortality rates. Hum Nat 17:74–97. doi:10.1007/s12110-006-1021-z

Kuroda T. 2018. A system for the real-time tracking of operant behavior as an application of 3D camera. Journal of the Experimental Analysis of Behavior 110:522–544. doi:10.1002/jeab.471

Liiver K, Imbeault S, Školnaja M, Kaart T, Kanarik M, Laugus K, De Wettinck J, Pulver A, Shimmo R, Harro J. 2023. Active vs passive novelty-related strategies: Sex differences in exploratory behaviour and monoaminergic systems. Behavioural Brain Research 441:114297. doi:10.1016/j.bbr.2023.114297

Lillesaar C. 2011. The serotonergic system in fish. Journal of Chemical Neuroanatomy 41:294–308. doi:10.1016/j.jchemneu.2011.05.009

Lillesaar C, Stigloher C, Tannhäuser B, Wullimann MF, Bally-Cuif L. 2009. Axonal projections originating from raphe serotonergic neurons in the developing and adult zebrafish, Danio rerio, using transgenics to visualize raphe-specific pet1 expression. Journal of Comparative Neurology 512:158–182. doi:10.1002/cne.21887

Lillesaar C, Tannhäuser B, Stigloher C, Kremmer E, Bally-Cuif L. 2007. The serotonergic phenotype is acquired by converging genetic mechanisms within the zebrafish central nervous system. Developmental Dynamics 236:1072–1084. doi:10.1002/dvdy.21095

Lima-Maximino M, Pyterson MP, do Carmo Silva RX, Gomes GCV, Rocha SP, Herculano AM, Rosemberg DB, Maximino C. 2020. Phasic and tonic serotonin modulate alarm reactions and post-exposure behavior in zebrafish. Journal of Neurochemistry 153:495–509. doi:10.1111/jnc.14978

Luca RM, Gerlai R. 2012. In search of optimal fear inducing stimuli: Differential behavioral responses to computer animated images in zebrafish. Behavioural Brain Research 226:66–76. doi:10.1016/j.bbr.2011.09.001

Lucki I. 1998. The spectrum of behaviors influenced by serotonin. Biological Psychiatry 44:151–162. doi:10.1016/S0006-3223(98)00139-5

Mathis A, Mamidanna P, Cury KM, Abe T, Murthy VN, Mathis MW, Bethge M. 2018. DeepLabCut: markerless pose estimation of user-defined body parts with deep learning. Nature Neuroscience 21:1281–1289. doi:10.1038/s41593-018-0209-y

Maximino C, P. Costa B, G. Lima M. 2016. A Review of Monoaminergic Neuropsychopharmacology in Zebrafish, 6 Years Later: Towards Paradoxes and their Solution. Current Psychopharmacology 5:96–138.

Maximino C, Puty B, Benzecry R, Araújo J, Lima MG, de Jesus Oliveira Batista E, Renata de Matos Oliveira K, Crespo-Lopez ME, Herculano AM. 2013. Role of serotonin in zebrafish (Danio rerio) anxiety: Relationship with serotonin levels and effect of buspirone, WAY 100635, SB 224289, fluoxetine and para-chlorophenylalanine (pCPA) in two behavioral models. Neuropharmacology 71:83–97. doi:10.1016/j.neuropharm.2013.03.006

McMillan StephanieC, Géraudie J, Akimenko M-A. 2015. Pectoral Fin Breeding Tubercle Clusters: A Method to Determine Zebrafish Sex. Zebrafish 12:121–123. doi:10.1089/zeb.2014.1060

Meager JJ, Fernö A, Skjæraasen JE, Järvi T, Rodewald P, Sverdrup G, Winberg S, Mayer I. 2012. Multidimensionality of behavioural phenotypes in Atlantic cod, Gadus morhua. Physiology & Behavior 106:462–470. doi:10.1016/j.physbeh.2012.03.010

Meyer A, Van de Peer Y. 2005. From 2R to 3R: evidence for a fish-specific genome duplication (FSGD). BioEssays 27:937–945. doi:10.1002/bies.20293

Moiron M, Laskowski KL, Niemelä PT. 2020. Individual differences in behaviour explain variation in survival: a meta-analysis. Ecology Letters 23:399–408. doi:10.1111/ele.13438

Montag C, Reuter M. 2014. Disentangling the molecular genetic basis of personality: From monoamines to neuropeptides. Neuroscience & Biobehavioral Reviews 43:228–239. doi:10.1016/j.neubiorev.2014.04.006

Näslund J, Studer E, Pettersson R, Hagsäter M, Nilsson S, Nissbrandt H, Eriksson E. 2015. Differences in Anxiety-Like Behavior within a Batch of Wistar Rats Are Associated with Differences in Serotonergic Transmission, Enhanced by Acute SRI Administration, and Abolished By Serotonin Depletion. International Journal of Neuropsychopharmacology 18:pyv018. doi:10.1093/ijnp/pyv018

Nichols DE, Nichols CD. 2008. Serotonin Receptors. Chem Rev 108:1614–1641. doi:10.1021/cr078224o

Norton WHJ, Folchert A, Bally-Cuif L. 2008. Comparative analysis of serotonin receptor (HTR1A/HTR1B families) and transporter (slc6a4a/b) gene expression in the zebrafish brain. Journal of Comparative Neurology 511:521–542. doi:10.1002/cne.21831

Nowicki M, Tran S, Muraleetharan A, Markovic S, Gerlai R. 2014. Serotonin antagonists induce anxiolytic and anxiogenic-like behavior in zebrafish in a receptor-subtype dependent manner. Pharmacology Biochemistry and Behavior 126:170–180. doi:10.1016/j.pbb.2014.09.022

Ochocki AJ, Kenney JW. 2023. A gelatin-based feed for precise and non-invasive drug delivery to adult zebrafish. Journal of Experimental Biology 226:jeb245186. doi:10.1242/jeb.245186

Press WH, Teukolsky SA. 1990. Savitzky-Golay smoothing filters. Computers in Physics 4:669–672.

R Core Team. 2016. R: A Language and Environment for Statistical Computing. R Foundation for Statistical Computing 1:409. doi:10.1007/978-3-540-74686-7

Rajput N, Parikh K, Kenney JW. 2022. Beyond bold versus shy: Zebrafish exploratory behavior falls into several behavioral clusters and is influenced by strain and sex. Biology Open 11:bio059443. doi:10.1242/bio.059443

Réale D, Reader SM, Sol D, McDougall PT, Dingemanse NJ. 2007. Integrating animal temperament within ecology and evolution. Biological Reviews 82:291–318. doi:10.1111/j.1469-185X.2007.00010.x

Ren G, Li S, Zhong H, Lin S. 2013. Zebrafish Tyrosine Hydroxylase 2 Gene Encodes Tryptophan Hydroxylase *. Journal of Biological Chemistry 288:22451–22459. doi:10.1074/jbc.M113.485227

Sackerman J, Donegan JJ, Cunningham CS, Nguyen NN, Lawless K, Long A, Benno RH, Gould GG. 2010. Zebrafish Behavior in Novel Environments: Effects of Acute Exposure to Anxiolytic Compounds and Choice of Danio rerio Line. International journal of comparative psychology 23:43–61.

Sánchez C, Bøgesø KP, Ebert B, Reines EH, Braestrup C. 2004. Escitalopram versus citalopram: the surprising role of the R-enantiomer. Psychopharmacology 174:163–176. doi:10.1007/s00213-004-1865-z

Schultz W. 2010. Dopamine signals for reward value and risk: basic and recent data. Behavioral and Brain Functions 6:24. doi:10.1186/1744-9081-6-24

Setini A, Pierucci F, Senatori O, Nicotra A. 2005. Molecular characterization of monoamine oxidase in zebrafish (Danio rerio). Comparative Biochemistry and Physiology Part B: Biochemistry and Molecular Biology 140:153–161. doi:10.1016/j.cbpc.2004.10.002

Spence R, Fatema MK, Reichard M, Huq KA, Wahab MA, Ahmed ZF, Smith C. 2006. The distribution and habitat preferences of the zebrafish in Bangladesh. Journal of Fish Biology 69:1435–1448. doi:10.1111/j.1095-8649.2006.01206.x

Teraoka H, Russell C, Regan J, Chandrasekhar A, Concha ML, Yokoyama R, Higashi K, Take-uchi M, Dong W, Hiraga T, Holder N, Wilson SW. 2004. Hedgehog and Fgf signaling pathways regulate the development of tphR-expressing serotonergic raphe neurons in zebrafish embryos. Journal of Neurobiology 60:275–288. doi:10.1002/neu.20023

Tran S, Nowicki M, Muraleetharan A, Chatterjee D, Gerlai R. 2016. Neurochemical factors underlying individual differences in locomotor activity and anxiety-like behavioral responses in zebrafish. Progress in Neuro-Psychopharmacology and Biological Psychiatry 65:25–33. doi:10.1016/j.pnpbp.2015.08.009

Verheij MMM, Veenvliet JV, Groot Kormelink T, Steenhof M, Cools AR. 2009. Individual differences in the sensitivity to serotonergic drugs: a pharmacobehavioural approach using rats selected on the basis of their response to novelty. Psychopharmacology 205:441–455. doi:10.1007/s00213-009-1552-1

Vormfelde SV, Hoell I, Tzvetkov M, Jamrozinski K, Sehrt D, Brockmöller J, Leibing E. 2006. Anxiety- and novelty seeking-related personality traits and serotonin transporter gene polymorphisms. Journal of Psychiatric Research 40:568–576. doi:10.1016/j.jpsychires.2005.10.002

Walther DJ, Bader M. 2003. A unique central tryptophan hydroxylase isoform. Biochemical Pharmacology 66:1673–1680. doi:10.1016/S0006-2952(03)00556-2

Wickham H. 2015. Elegant Graphics for Data Analysis, Media. Springer. doi:10.1007/978-0-387-98141-3

Wilson DS, Clark AB, Coleman K, Dearstyne T. 1994. Shyness and boldness in humans and other animals. Trends in Ecology & Evolution 9:442–446. doi:10.1016/0169-5347(94)90134-1

Wilson M, Daly M. 1985. Competitiveness, risk taking, and violence: the young male syndrome. Ethology and Sociobiology 6:59–73. doi:10.1016/0162-3095(85)90041-X

Winberg S, Thörnqvist P-O. 2016. Role of brain serotonin in modulating fish behavior. Current Zoology 62:317–323. doi:10.1093/cz/zow037

