## Supplemental Figures for "Individual differences in the boldness of female zebrafish are associated with alterations in serotonin function"

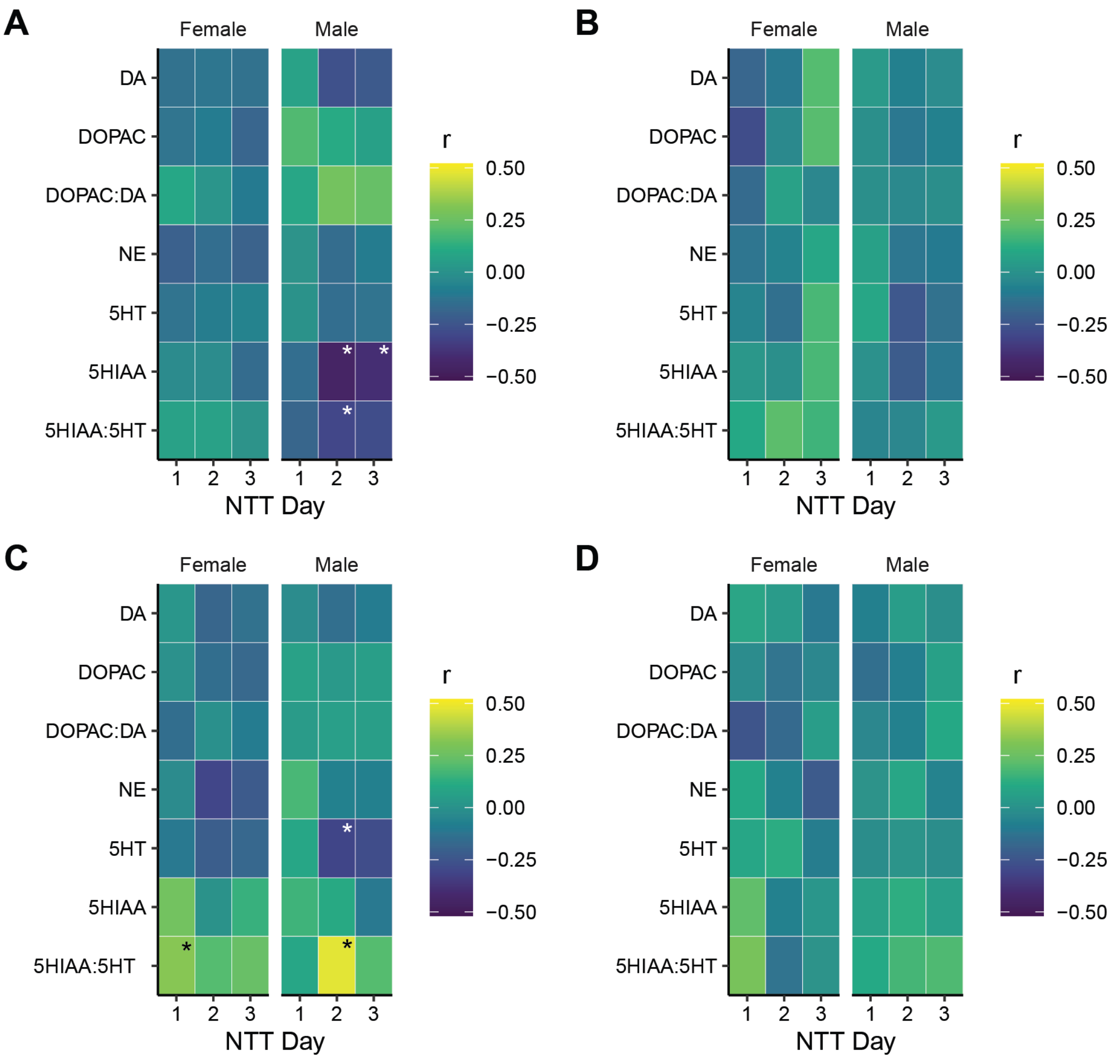


**Figure S1.** Pearson's correlations between individual behavioral parameters and brain chemistry in male and female zebrafish. A) distance from bottom, B) distance from center, C) distance travelled, and D) percent tank explored. * - $P < 0.05$; n's same as in figure 1.

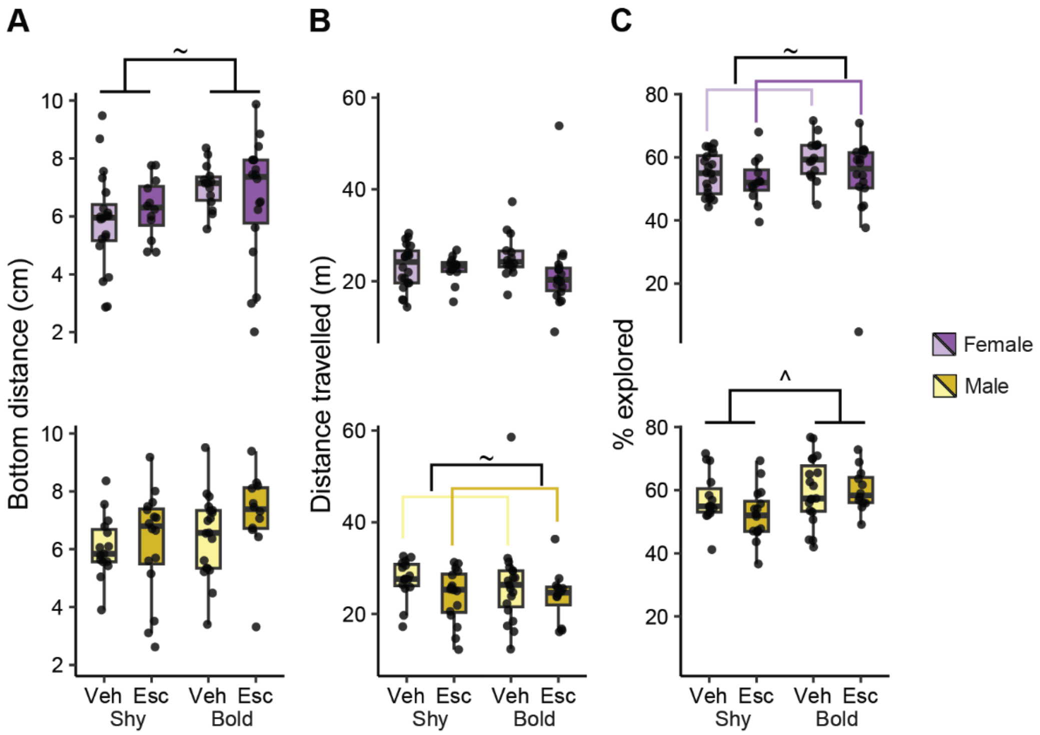


**Figure S2.** The effect of escitalopram on behavior in bold and shy fish. Experimental setup is the same as in Figure 3A. A) Bottom distance, B) distance travelled, and C) percent explored. Boxplot center is the median, hinges are interquartile ranges, and whiskers are the hinge ± 1.5 times the interquartile range. ~ - $P < 0.10$, ^ - $P < 0.05$ from two way ANOVA; n's same as in Figure 3.

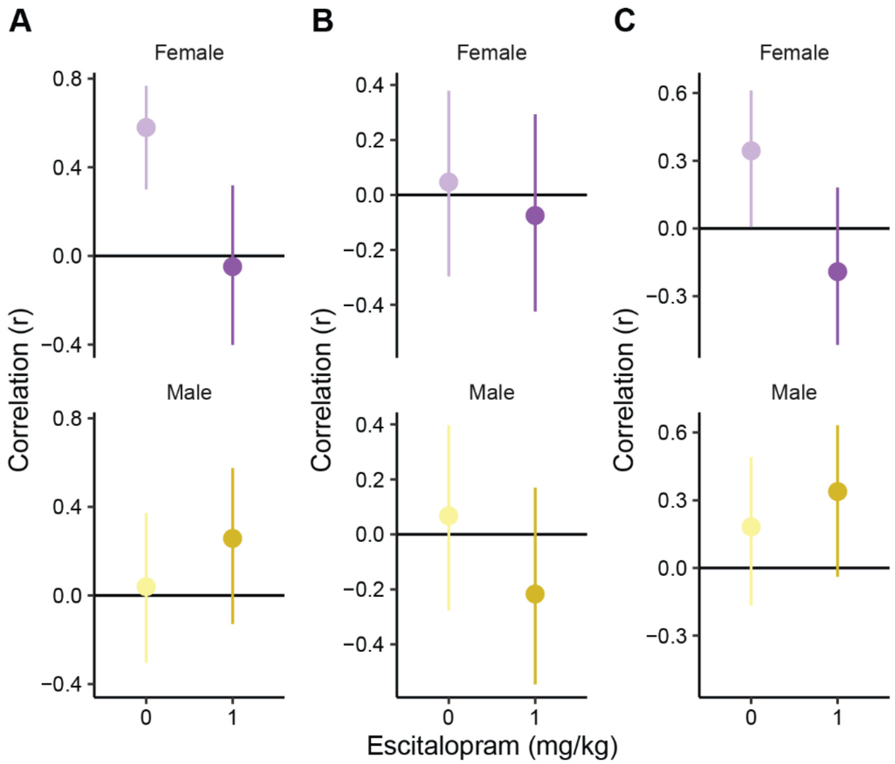


**Figure S3.** Pearson's correlations between boldness index on day 1 and day 2 behaviors of A) bottom distance, B) distance travelled, and C) percent explored. Experimental design same as in figure 3A. Error bars are 95% confidence intervals.
